# *TAS2R10* Suppresses ABCG2 Transporter-Associated Chemoresistance and Enhances Cisplatin Sensitivity in Head and Neck Squamous Cell Carcinoma

**DOI:** 10.64898/2026.07.26.740768

**Authors:** Lily Huang, Sarah M. Sywanycz, Priyanka Sahu, Leilei Hao, Kyle Polen, Gavin Turner, Zoey A. Miller, Robert J. Lee, Ryan M. Carey

## Abstract

Cisplatin resistance remains a major barrier in head and neck squamous cell carcinoma (HNSCC) treatment. ATP-binding cassette (ABC) transporters contribute to chemoresistance by limiting intracellular drug accumulation. Bitter taste receptor 10 (T2R10) has been implicated in ABC transporter regulation, but its role in HNSCC remains undefined.

HNSCC cell lines were treated with T2R10-agonist caffeine (100 or 200 μM), cisplatin, or a combination, and viability was assessed by crystal violet assay. T2R10 promoter activity and expression following caffeine exposure were evaluated using a promoter-driven mCherry reporter and RT-qPCR. ABC transporter expression was measured after caffeine treatment and T2R10 gene (*TAS2R10*) knockdown or overexpression. Associations between tumor *TAS2R10* expression and survival were assessed using TCGA data through GEPIA2.

Caffeine enhanced the cisplatin-associated reduction in viability in a cell line- and concentration-dependent manner, with the strongest effect seen in UM-SCC47. A significant effect was observed in FaDu at 200 μM of caffeine, and minimal response in RPMI 2650. RPMI 2650 cells and FaDu cells exhibited lower baseline *TAS2R10* expression and RPMI 2650 cells did not demonstrate enhanced cisplatin sensitivity following caffeine treatment. Caffeine treatment increased *TAS2R10* promoter activity and expression and was associated with decreased *ABCG2* expression. *TAS2R10* knockdown increased *ABCG2* and *ABCF1* expression, whereas *TAS2R10* overexpression reduced *ABCG2* and *ABCC1* expression. High tumor *TAS2R10* expression was associated with improved disease-free survival (log-rank p=0.0071; HR=0.61) but not overall survival.

Caffeine enhances cisplatin sensitivity in selected HNSCC models. Caffeine exposure is associated with increased *TAS2R10* expression and reduced expression of chemoresistance-associated transporters, particularly ABCG2.

## Introduction

Bitter taste receptors (T2Rs) are G protein-coupled receptors (GPCRs) classically involved in bitter compound detection on the tongue but are also expressed in extraoral tissues where they regulate diverse physiologic and pathologic processes.^1–3^ Increasing evidence supports a role for T2Rs in malignancies including head and neck squamous cell carcinoma (HNSCC). ^4–14^ In HNSCC, activation of T2R14 by lidocaine and T2R5 by phendione has been shown to reduce cell viability and promote apoptosis through mitochondrial depolarization, reactive oxygen species generation, and caspase activation.^6, 12, 14^ Increased expression of some T2R genes (known as *TAS2R*s) in patient tumors has also been associated with improved overall survival in TCGA cohorts, underscoring the potential relevance of these receptors.^6, 15^ Together, these studies suggest that T2Rs regulate pathways involved in tumor cell survival, apoptosis, and sensitivity to anticancer therapies in HNSCC, raising the possibility that T2R signaling may also influence sensitivity to conventional cytotoxic therapies.

Because cisplatin-based chemotherapy remains a cornerstone of treatment for locally advanced and recurrent or metastatic head and neck squamous cell carcinoma (HNSCC)^16, 17^, identifying mechanisms that regulate treatment sensitivity has substantial clinical importance. However, intrinsic and acquired resistance to cisplatin remain a major clinical obstacle that contributes to treatment failure, recurrence, and poor survival outcomes.^18, 19^ Mechanisms of cisplatin resistance include enhanced DNA repair, apoptosis evasion, and altered drug transport.^18, 19^ Among these, increased activity of ATP-binding cassette (ABC) transporters including ABCG2 reduce intracellular drug accumulation and contribute to chemoresistance in multiple solid tumors, including HNSCC.^19–21^ Despite extensive investigation, clinically effective ABC transporter inhibitors for cancer therapy have remained elusive. Accordingly, identifying upstream signaling pathways that regulate ABC transporter expression may represent an alternative strategy to overcome chemoresistance.^21^

Among T2Rs, T2R10 has recently been implicated in regulation of cancer-associated signaling pathways. Caffeine, a naturally occurring methylxanthine and known T2R10 agonist, therefore represents a clinically relevant means of interrogating T2R10 signaling. Epidemiologic studies have suggested that caffeinated coffee consumption may be associated with reduced risk of head and neck cancers.^22–24^ Previous studies demonstrated that caffeine-mediated T2R10 activation modulates Akt phosphorylation and ABC transporter expression in pancreatic adenocarcinoma.^1, 25, 26^ These findings raise the possibility that T2R10 activation may similarly influence therapeutic response in HNSCC through regulation of ABC transporter expression.

Whether caffeine-associated T2R10 signaling modulates cisplatin sensitivity in HNSCC remains unknown. Given the established role of ABC transporters in HNSCC chemoresistance and evidence that T2R10 signaling regulates transporter expression, we investigated whether caffeine enhances cisplatin cytotoxicity by modulating ABC transporter expression in HNSCC cell lines. We also evaluated the prognostic significance of T2R10 gene (*TAS2R10*) expression in HNSCC using The Cancer Genome Atlas (TCGA). We hypothesized that caffeine enhances cisplatin sensitivity by suppressing ABC transporter expression through T2R10-associated signaling.

## Materials and Methods

### Cell Culture

Human HNSCC cell lines SCC47 (lateral tongue, HPV positive) (UM-SCC-47; Millipore SCC071, RRID:CVCL_7759), FaDu (hypopharynx, HPV negative) (ATCC HTB-43, RRID:CVCL_1218), RPMI 2650 (nasal septum, HPV negative) (ATCC CCL-30, RRID:CVCL_1664), were obtained from MilliporeSigma (St. Louis, MO USA) or ATCC (Manassas, VA, USA). Cells were cultured in high glucose DMEM (Corning; Glendale, AZ, USA) supplemented with 10% FBS (Genesee Scientific; El Cajon, CA, USA), 1% penicillin/streptomycin mix (P/S) (Gibco; Gaithersburg, MD, USA), and 1% nonessential amino acids (NEAA)(Gibco).

### Reverse Transcription Quantitative PCR (RT-qPCR)

Cells were lysed and resuspended in TRIzol (Thermo Fisher Scientific), and RNA was extracted using a Direct-zol RNA kit (Zymo Research). Quantitative reverse transcription (qPCR) was performed using the High-Capacity cDNA Reverse Transcription Kit (Thermo Fisher Scientific), and *TAS2R10* expression was quantified using TaqMan qPCR assays. Housekeeping gene ubiquitin C (UBC) was selected for normalization due to its stable expression in cancer cells. ^27^

### ABC Transporter Expression

UM-SCC47 cells were treated with caffeine (100 μM or 200 μM) for 72 hours with re-dosing every 12 hours. Total RNA was isolated using TRIzol and purified using the Direct-zol RNA kit (Zymo Research). Reverse transcription was performed using the High-Capacity cDNA Reverse Transcription Kit (Thermo Fisher Scientific). Expression of *ABCG2*, *ABCC1*, *ABCF1*, and *TAP2* was quantified using TaqMan RT-qPCR assays and normalized to the housekeeping gene *UBC*. Relative gene expression was normalized to untreated controls.

To evaluate the effects of caffeine in patient-derived tissue, similar experiments were performed using ex vivo tumor slices generated from HPV-positive oropharyngeal squamous cell carcinoma specimens collected under a protocol approved by the University of Pennsylvania Institutional Review Board (IRB #417200), as previously described.^14^ Tumor specimens were sectioned into 300-μm-thick slices using a Precisionary Compresstome VF-510-0Z vibratome with the oscillation and speed settings set to 6 and 3, respectively. Tumor slices were cultured in control media or media containing 100 or 200 μM caffeine. RNA was subsequently extracted, and qPCR was performed as described above.

### TAS2R10 Promoter Assay

To evaluate the effect of caffeine on *TAS2R10* promoter activity, cells were transfected with an mCherry reporter construct driven by the *TAS2R10* promoter. Following transfection, cells were treated with 200 μM caffeine. mCherry fluorescence was assessed following treatment as a measure of *TAS2R10* promoter activity. In parallel, *TAS2R10* expression following caffeine exposure was evaluated by RT-qPCR normalized to *UBC*.

### Knockdown and Overexpression of TAS2R10

UM-SCC47 cells were transfected with *siTAS2R10* or non-targeting siScramble control using RNAiMAX lipofectamine reagent (Thermo Fisher Scientific) and Opti-MEM (Gibco). Suspended transfections were performed on day 1 with an additional transfection on day 3. Cells were collected for downstream assays following transfection. Knockdown efficiency was validated by RT-qPCR, demonstrating reduced *TAS2R10* expression compared to siScramble controls.

UM-SCC47 cells were transfected with a pcDNA3.1(+) vector containing full-length human *TAS2R10* using Lipofectamine 3000 and P3000 reagent (Thermo Fisher Scientific) according to the manufacturer’s instructions. Cells were collected 24–48 hours post-transfection for downstream assays. Overexpression efficiency was assessed by RT-qPCR, which demonstrated marked upregulation of *TAS2R10* mRNA compared to empty vector controls.

### The Cancer Genome Atlas (TCGA) Analysis

Kaplan–Meier survival analysis of *TAS2R10* expression in head and neck squamous cell carcinoma was performed using the GEPIA2 web server, which utilizes RNA sequencing data from The Cancer Genome Atlas (TCGA) and GTEx databases. Patients were stratified into high and low *TAS2R10* expression groups using the default GEPIA2 settings. Overall survival and disease-free survival curves were generated using the GEPIA2 survival analysis module, and statistical significance was assessed using the log-rank test.

### Cell Viability

Cells were treated with caffeine and cisplatin in high-glucose DMEM supplemented with 10% FBS, 1% penicillin/streptomycin, and 1% nonessential amino acids at 37° C in a humidified incubator at 5% CO_2_. Sublethal caffeine dosing was first established through caffeine-only crystal violet assays, and concentrations of 100 μM and 200 μM were selected. Cisplatin concentrations were selected based on IC_50_ determination. UM-SCC-47 was treated with 10 μM cisplatin, while FaDu and RPMI 2650 were treated with 5 μM cisplatin. Cells were treated with caffeine every 12 hours over a total treatment period of 72 hours. After 24 hours of caffeine exposure, cisplatin was added once at the indicated IC50 concentration. At the completion of the treatment period, remaining adherent cells were stained with 0.1% crystal violet, washed, dried, and dissolved in 30% acetic acid. Absorbance at 590 nm was measured using a Tecan plate reader (Tecan Spark 10M; Mannedorf, Switzerland) and normalized to untreated controls.

### Statistical Analyses

Statistical analyses were conducted using GraphPad Prism (San Diego, CA, USA). T-tests were used for two-group comparisons, while one-way ANOVA was applied for comparisons across multiple groups. Sample sizes were selected based on prior publications demonstrating sufficient power to detect treatment effects in similar models; no formal power analysis was performed. Statistical tests used are described in the figure legends, with significance indicated as follows: P < 0.05 (*), P < 0.01 (**), P < 0.001 (***), P < 0.0001 (****). Error bars represent mean ± SEM.

### Data Availability

All data generated in this study are shown in the manuscript. The raw data used to generate graphs and figures are available from the corresponding author upon reasonable request.

## Results

### TAS2R10 is expressed in HNSCC and is associated with favorable clinical outcomes

To determine whether the *TAS2R10* gene that encodes T2R10 is expressed in HNSCC cell lines, *TAS2R10* expression was quantified by RT-qPCR and normalized to housekeeping gene ubiquitin C (UBC). Relative *TAS2R10* expression differed across the three cell lines, with higher expression observed in UM-SCC47 compared to FaDu and RPMI 2650 (Figure 1).

**Figure 1.**
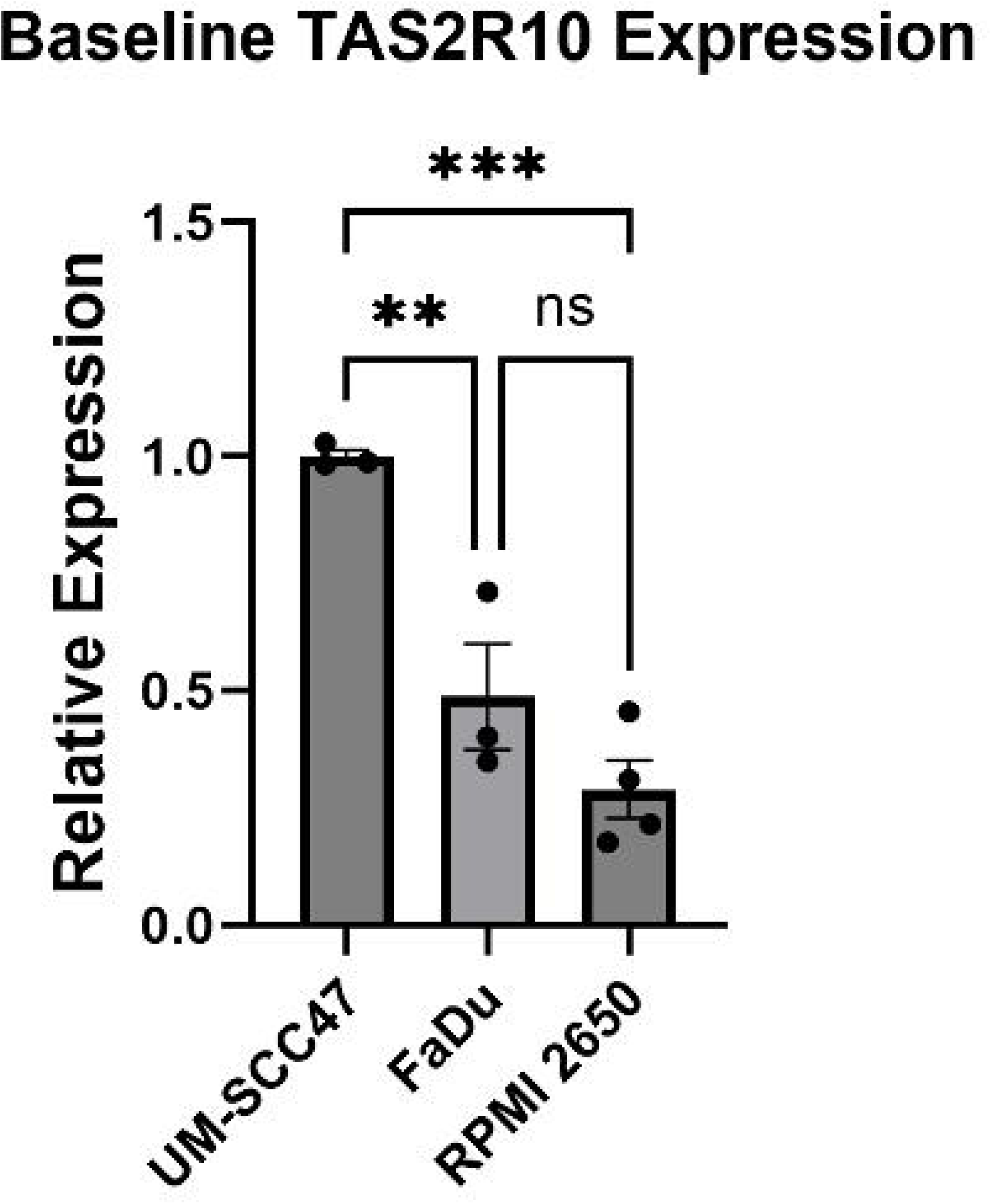
TAS2R10 is expressed in HNSCC. Relative expression of *TAS2R10* in cell lines ΜM-SCC47, FaDu, and RPMI 2650. Expression of each TAS2R mRNA was normalized to ubiquitin C (UBC) housekeeping gene. Significance by 1-way ANOVA with Dunnett’s multiple comparison posttest. *p < 0.05, **p < 0.01, ***p < 0.001; ns, no statistical significance.

To evaluate the relationship of T2R10 and clinical outcomes in HNSCC, we evaluated disease free survival in patients with tumors expressing high versus low levels of *TAS2R10*. Kaplan-Meier analysis using TCGA datasets demonstrated that high *TAS2R10* expression was associated with significantly improved disease-free survival compared to low *TAS2R10* levels (log-rank p = 0.0071; HR = 0.61) (Figure 2A). In contrast, *TAS2R10* expression was not associated with overall survival (Figure 2B). These findings suggest that increased *TAS2R10* expression is associated with favorable disease-free survival in HNSCC.

**Figure 2.**
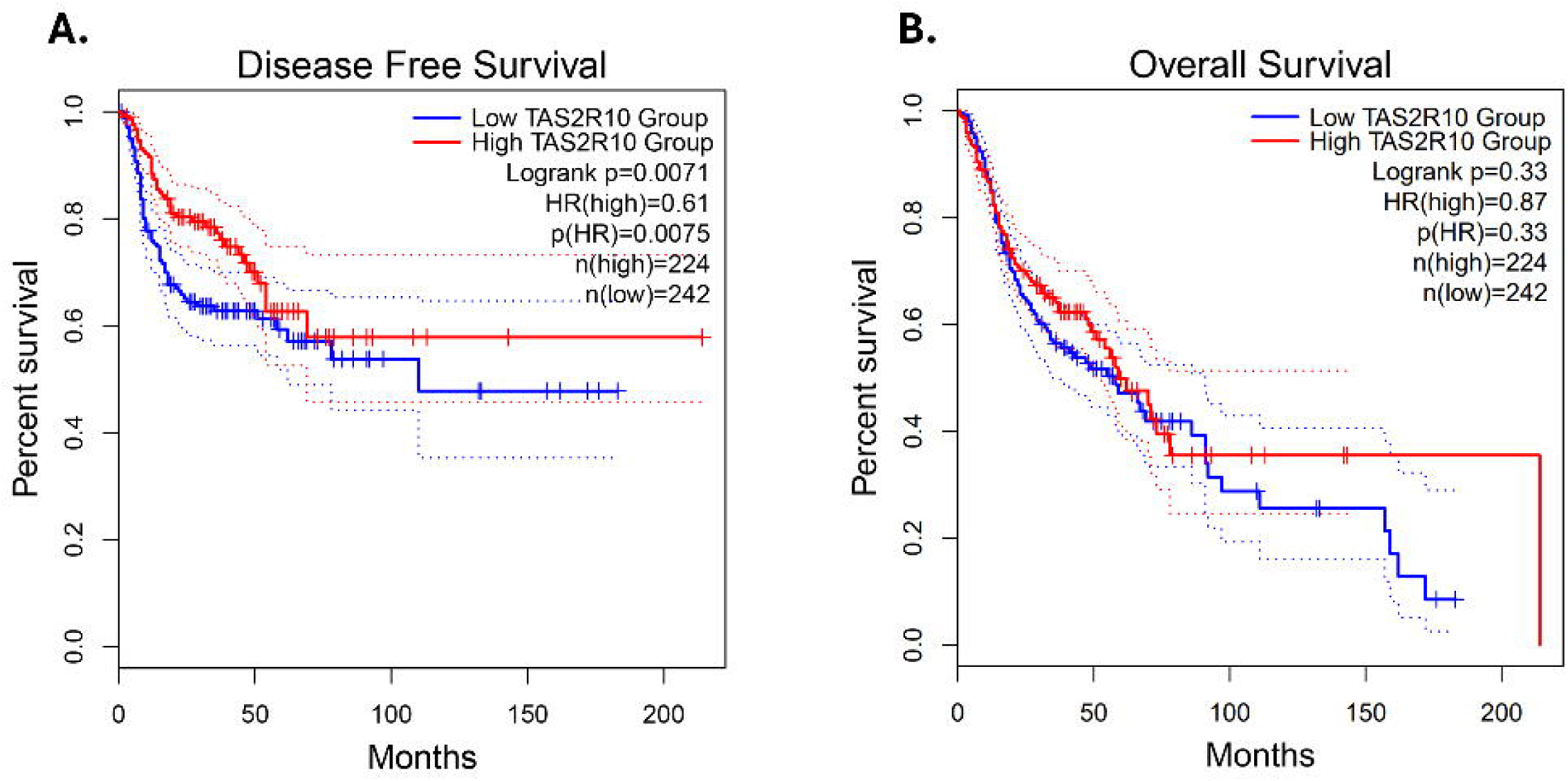
TAS2R10 expression and survival in HNSCC. Kaplan–Meier **(A)** disease-free survival (HR = 0.61, p = 0.0071) and **(B)** overall survival (HR = 0.87, p = 0.33) analysis of patients with head and neck squamous cell carcinoma stratified by high versus low TAS2R10 expression using TCGA datasets accessed through the GEPIA2 web server. Patients were dichotomized into high and low expression groups using a 50% cutoff (upper 50% vs lower 50%). Statistical significance was assessed using the log-rank test.

### TAS2R10 expression is inversely associated with ABCG2 transporter expression

Given the association between *TAS2R10* expression and improved disease-free survival, we next investigated whether T2R10 modulates pathways associated with chemoresistance. Because ABC transporters contribute to drug resistance by limiting intracellular drug accumulation, we examined whether experimental manipulation of *TAS2R10* altered ABC transporter gene expression in UM-SCC47 cells. To assess the effect of *TAS2R10* loss, UM-SCC47 cells were transfected with si*TAS2R10* or non-targeting siScramble control. Following *TAS2R10* knockdown, *ABCG2* expression increased (Figure 3A), suggesting that reduced *TAS2R10* expression may promote an elevated ABCG2 state associated with chemoresistance.

**Figure 3.**
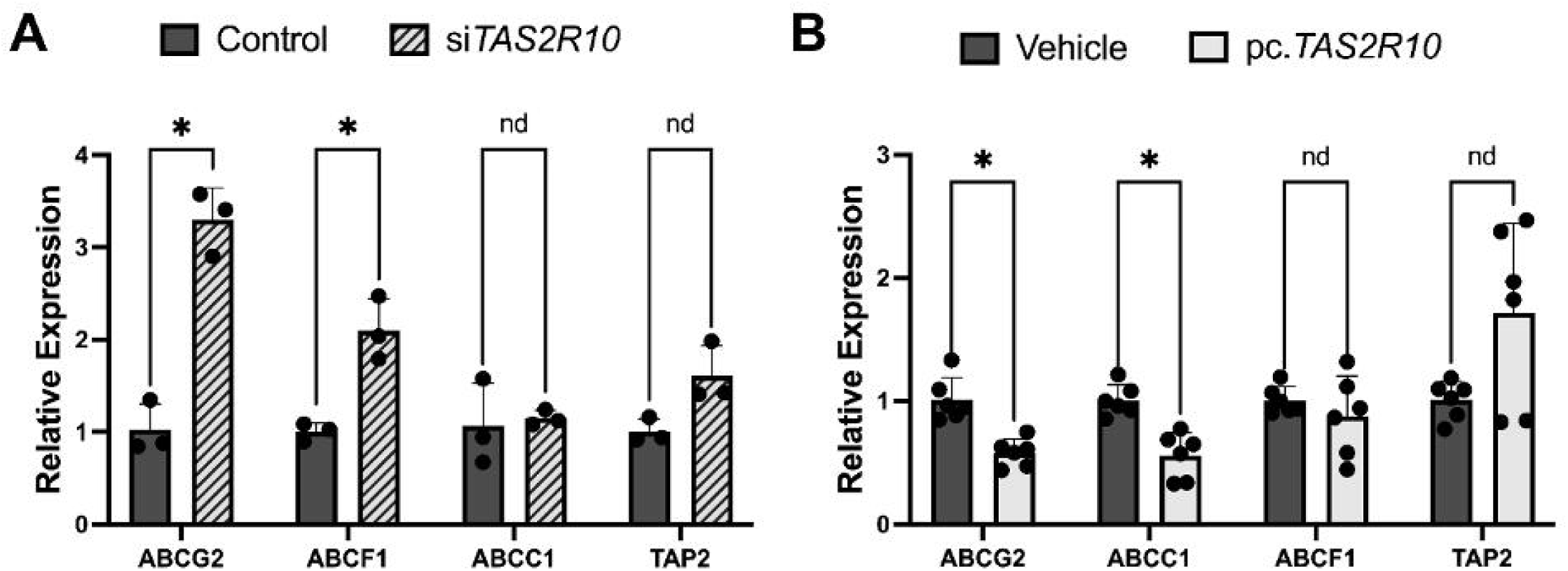
Modulation of TAS2R10 alters ABC Transporter expression. **A)** Relative expression of ABCG2, ABCF1, ABCC1, and TAP2 following *TAS2R10* knockdown compared with control. **B)** Relative expression of ABCG2, ABCC1, ABCF1, and TAP2 following *TAS2R10* overexpression compared with vehicle control. Expression was measured by qRT-PCR and normalized to UBC. Statistical significance was determined by unpaired *t*-test for two-group comparisons. *p < 0.05, **p < 0.01, ***p < 0.001; ns, no statistical significance.

To determine whether increased *TAS2R10* expression produced the reciprocal effect, UM-SCC47 cells were transfected with a full-length *TAS2R10* overexpression construct or empty vector control. *TAS2R10* overexpression was confirmed by RT-qPCR and was associated with reduced expression of *ABCG2* (Figure 3B). Among the ABC transporter genes evaluated, *ABCG2* demonstrated the most consistent reciprocal relationship with *TAS2R10* modulation. Together, these gain- and loss-of-function experiments support an inverse relationship between *TAS2R10* expression and ABCG2 expression. These findings suggest that *TAS2R10* may function as an upstream regulator of ABCG2 transporter-associated chemoresistance pathways in HNSCC cells.

### Caffeine increases TAS2R10 promoter activity and TAS2R10 expression

Having established a relationship between *TAS2R10* expression and ABC transporter regulation, we next asked whether caffeine, a T2R10 agonist, modulates *TAS2R10* expression in HNSCC cells. UM-SCC47 cells were transfected with a *TAS2R10* promoter-driven mCherry reporter construct and treated with caffeine. Caffeine exposure increased mCherry fluorescence compared with control conditions, indicating increased *TAS2R10* promoter activity (Figure 4A).

**Figure 4.**
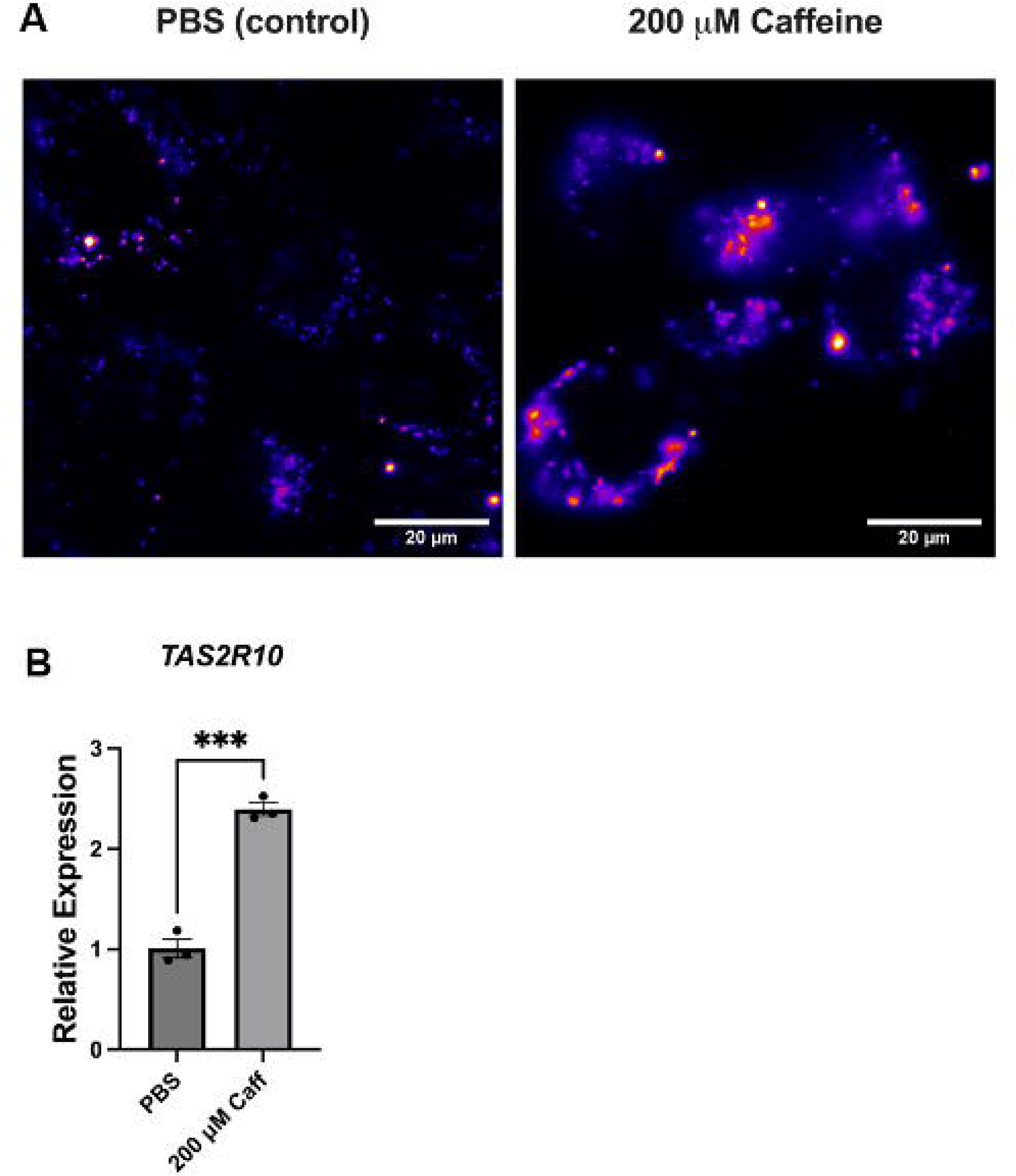
Caffeine increases TAS2R10 promoter activity and TAS2R10 expression in SCC47 cells. **A)** Representative fluorescence microscopy images of SCC47 cells expressing a TAS2R10 promoter-driven mCherry reporter under PBS control conditions or following treatment with 200 µM caffeine. Increased mCherry fluorescence after caffeine treatment is consistent with enhanced TAS2R10 promoter activity. **B)** Relative TAS2R10 mRNA expression in SCC47 cells following treatment with 200 µM caffeine compared with PBS control. Expression was measured by qRT-PCR, normalized to UBC, and expressed relative to PBS-treated cells. Statistical significance determined by unpaired t-test. *p < 0.05, **p < 0.01, ***p < 0.001; ns, no statistical significance.

Consistent with the promoter reporter findings, RT-qPCR demonstrated increased *TAS2R10* mRNA expression following caffeine treatment (Figure 4B). These findings demonstrate that caffeine increases *TAS2R10* promoter activity and mRNA expression in HNSCC cells.

### Caffeine consistently reduces ABCG2 expression

Because *TAS2R10* overexpression reduced *ABCG2* transporter gene expression and caffeine increased *TAS2R10* promoter activity and expression, we next evaluated whether caffeine treatment similarly altered *ABCG2* transporter expression. UM-SCC47 cells were treated with caffeine at 100 μM or 200 μM for 72 hours, with re-dosing every 12 hours. *ABCG2* transporter gene expression was then quantified by RT-qPCR and normalized to untreated controls. The same caffeine treatment was also applied to patient-derived tumor tissue.

Caffeine exposure significantly reduced *ABCG2* expression in both patient-derived tumor tissue and UM-SCC47, with the greatest reduction observed at 200 μM (Figure 5). The direction of this effect is consistent with the transporter changes observed following *TAS2R10* overexpression, supporting a model in which caffeine may reduce *ABCG2*-associated chemoresistance markers through *TAS2R10*-associated regulation.

**Figure 5.**
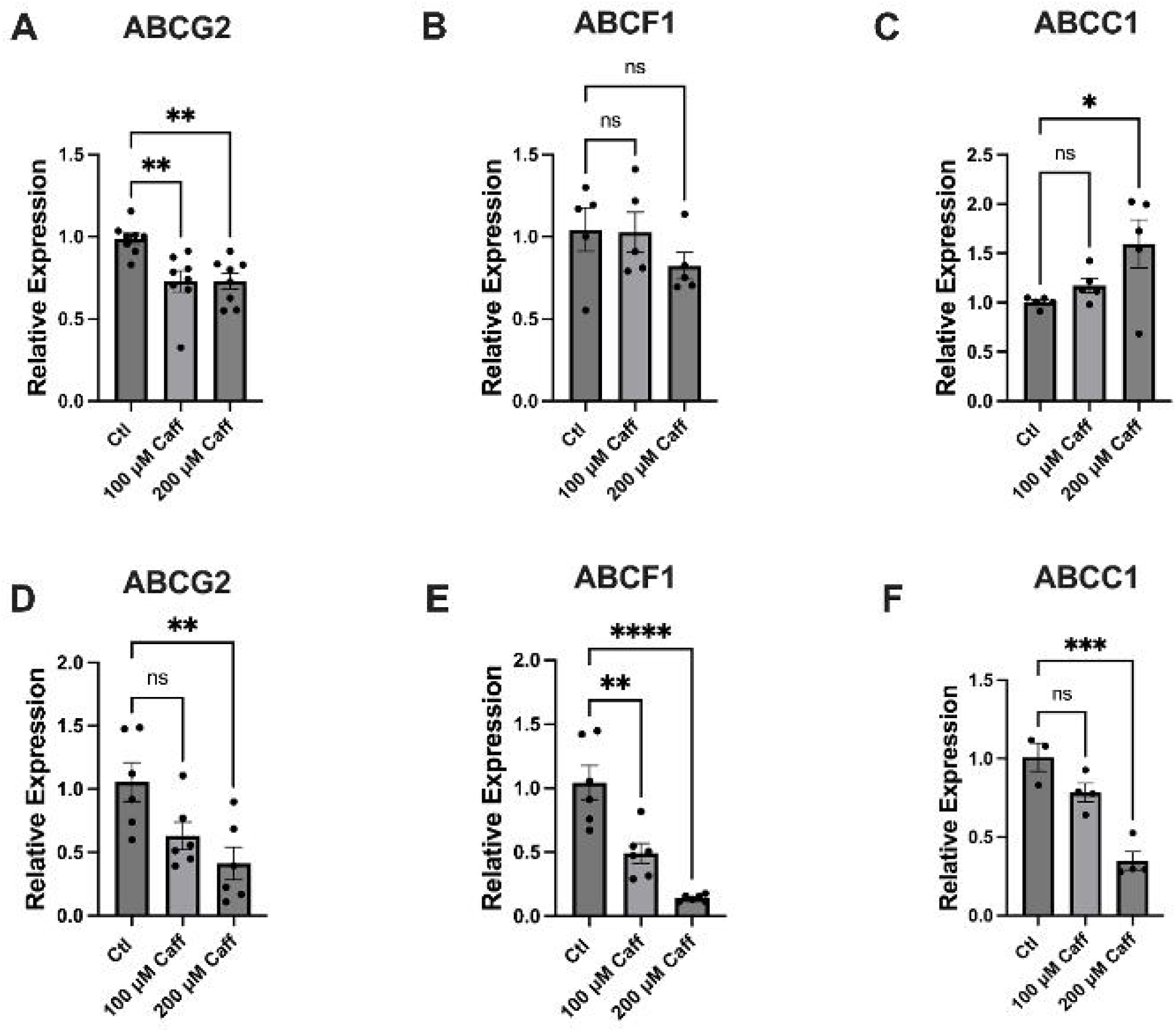
Caffeine treatment and modulation of *TAS2R10* alter ABC transporter expression. **A–C)** Relative mRNA expression of ABCG2, ABCF1, and ABCC1 in SCC47 cells following 72-hour treatment with control, 100 µM caffeine, or 200 µM caffeine. Expression was measured by qRT-PCR, normalized to *UBC*, and expressed relative to control-treated cells. **D–F)** Relative mRNA expression of ABCG2, ABCF1, and ABCC1 in tumor slice culture cells following 72-hour treatment with control, 100 µM caffeine, or 200 µM caffeine, measured and normalized as above. Statistical Significance determined by one-way ANOVA for caffeine treatment experiments with Dunnett’s multiple comparison posttest. *p < 0.05, **p < 0.01, ***p < 0.001; ns, no statistical significance.

Other ABC transporters demonstrated more variable responses to caffeine treatment. In UM-SCC47 cells, *ABCC1* expression increased after caffeine treatment, contrary to the tumor slice cultures where *ABCC1* decreased. *ABCF1* expression also differed between UM-SCC47 and tumor slice cultures following caffeine treatment with non-significant changes in the cell line but a significant decrease in tumor slice cultures. These findings suggest that caffeine-associated regulation of ABC transporter genes is not uniform across transporters or model systems. Therefore, subsequent interpretation focused on *ABCG2*, which showed the most consistent inverse association with caffeine treatment and *TAS2R10* activity. Because ABCG2 has been implicated in cisplatin resistance in HNSCC, these findings provide a potential mechanistic basis for the effects of caffeine on cisplatin sensitivity evaluated in the subsequent experiments.

### Caffeine enhances cisplatin sensitivity in HNSCC cell lines with higher TAS2R10 expression

Finally, we evaluated whether caffeine-associated changes in *TAS2R10* and *ABCG2* expression were accompanied by altered cisplatin sensitivity. Sublethal caffeine concentrations of 100 μM and 200 μM were selected based on viability assays with caffeine alone. HNSCC cells were treated with media control, caffeine alone, cisplatin alone, or caffeine in combination with cisplatin. UM-SCC47 cells were treated with 10 μM cisplatin, while FaDu and RPMI 2650 cells were treated with 5 μM cisplatin. Cell viability was assessed by crystal violet staining after 72 hours.

Caffeine alone had minimal or no significant effect on cell viability compared with media (vehicle) controls. However, caffeine enhanced cisplatin-associated reduction in cell viability in UM-SCC47 and FaDu cells (Figure 6). The strongest combination effect was observed in UM-SCC47 cells, which also demonstrated relatively high baseline *TAS2R10* expression. In contrast, RPMI 2650 cells, which expressed lower *TAS2R10*, did not show a clear enhancement of cisplatin sensitivity with caffeine treatment.

**Figure 6.**
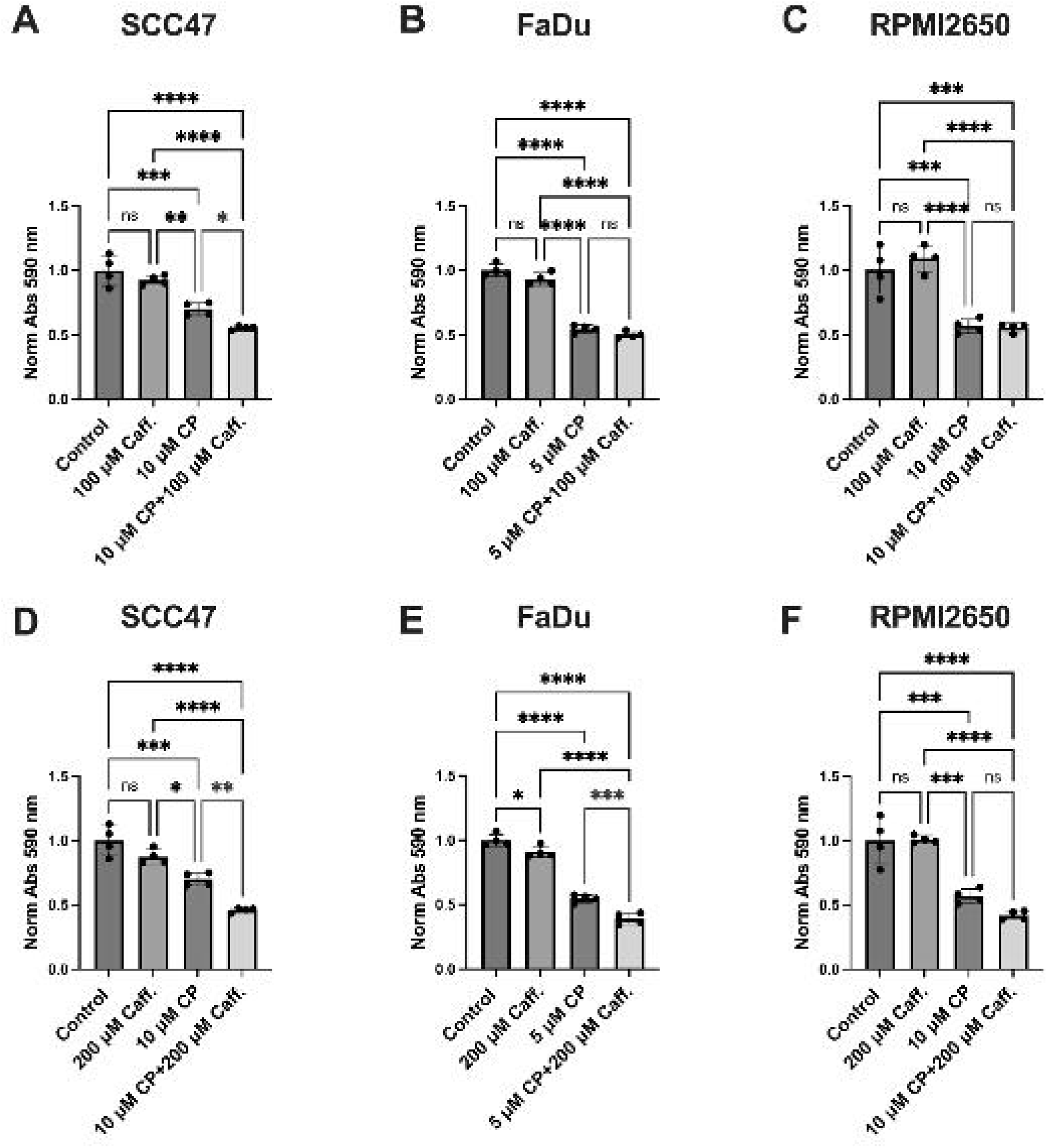
Caffeine may sensitize HNSCC cells to cisplatin-mediated decreases in cell viability. Crystal violet cell viability for **A)** UM-SCC47, **B)** FaDu, and **C)** RPMI 2650 treated with 10 μM cisplatin (CP) and 100 μM caffeine (Caff), and **D)** UM-SCC47 **E)** FaDu, and **F)** RPMI 2650 treated with 10 μM cisplatin and 200 μM caffeine. Significance by 1-way ANOVA with Dunnett’s multiple comparison posttest. *p < 0.05, **p < 0.01, ***p < 0.001; ns, no statistical significance.

These findings suggest that caffeine acts primarily as a chemosensitizing agent at the selected concentrations rather than as an independent cytotoxic treatment. The differential response across cell lines suggests that baseline *TAS2R10* expression may contribute to sensitivity to caffeine-associated cisplatin enhancement.

## Discussion

Our study demonstrates that caffeine sensitizes HNSCC cells to cisplatin and is associated with decreased expression of the *ABCG2* gene, which encodes a transporter implicated in chemoresistance. Caffeine exposure also increased *TAS2R10* promoter activity and *TAS2R10* expression, suggesting that caffeine may modulate T2R10-associated pathways through effects on receptor expression rather than acute receptor activation alone. In addition, bidirectional modulation of *TAS2R10* expression demonstrated an inverse relationship between *TAS2R10* and *ABCG2* expression, further supporting a link between T2R10 signaling and transporter-associated chemoresistance pathways. Given the established role of ABC transporters, particularly ABCG2, in HNSCC chemoresistance,^20, 28^ these findings suggest that T2R10-associated signaling may influence chemotherapy response in HNSCC. If confirmed mechanistically, modulation of T2R10 signaling could represent an alternative strategy to indirectly suppress ABCG2-associated chemoresistance, thereby avoiding many of the limitations encountered with direct ABC transporter inhibitors.

Previous studies in pancreatic adenocarcinoma demonstrated that caffeine-associated T2R10 signaling can suppress Akt phosphorylation and downregulate ABCG2 transporter expression.^1^ Similarly, in lung and colorectal cancer models, caffeine enhanced chemotherapy-induced cytotoxicity by modulating DNA damage response and apoptosis.^25, 29^ Together, these studies support the concept that caffeine may sensitize epithelial cancers to chemotherapy through multiple mechanisms, potentially including the modulation of *TAS2R10*-associated signaling and ABCG2 expression. ABCG2 was the most consistent transporter associated with caffeine treatment and *TAS2R10* modulation, whereas other ABC transporters demonstrated more variable responses. For example, *ABCC1* increased in UM-SCC47 cells following caffeine treatment but decreased in tumor slice cultures and following *TAS2R10* overexpression, suggesting transporter-specific and model-dependent regulation. These findings may reflect differences between isolated cell culture and the tumor microenvironment in patient-derived tumor slice cultures, as well as potential T2R10-independent effects of caffeine. Accordingly, ABCG2 represented the most consistent transporter-associated readout in this study, although additional functional studies will be needed to determine whether ABCG2 directly mediates caffeine-associated cisplatin sensitization.

RPMI 2650 cells did not exhibit enhanced cisplatin sensitivity following caffeine treatment while the effect in FaDu cells were only observed at 200 μM of caffeine. Notably, these cells also expressed lower baseline *TAS2R10* levels than UM-SCC47. Although multiple cell line-specific factors may contribute to this difference, these findings raise the possibility that baseline *TAS2R10* expression influences responsiveness to caffeine-associated signaling.

Prior studies in HNSCC have demonstrated that T2R activation promotes apoptosis through calcium signaling, mitochondrial depolarization, and reactive oxygen species generation.^6, 12^ Our group previously demonstrated that activation of T2R14 with lidocaine and T2R5 with phendione reduces HNSCC cell viability and promotes apoptosis.^12, 14^ Building on these findings, our group is conducting a phase I clinical trial investigating neoadjuvant intratumoral lidocaine in HNSCC.^30^ Beyond apoptosis, accumulating evidence suggests that T2Rs may regulate broader oncogenic and therapeutic response pathways in HNSCC and other solid tumors. For example, *TAS2R10* has been associated with suppression of aggressive tumor phenotypes in neuroblastoma models, while loss of T2R14 has been linked to increased mechanistic target of rapamycin (mTOR) signaling in HNSCC.^31, 32^ Across multiple studies, increased TAS2R expression has been associated with improved survival in HNSCC, which is consistent with our finding that high *TAS2R10* expression correlates with improved disease-free survival. Together, the current literature supports an emerging role of T2Rs as regulators of oncogenic signaling pathways and therapeutic response in HNSCC.

Emerging evidence suggests that the role of T2Rs in chemoresistance is both receptor and context dependent. Recent work in breast cancer demonstrated that intracellular T2Rs, particularly T2R14, can promote chemoresistance by upregulating ABCB1 through a Gα12/13–RhoA/ROCK–p38MAPK–NF-κB pathway.^33^ In these models, exposure to T2R ligands enhanced drug efflux and facilitated doxorubicin resistance. Additionally, pancreatic cancer models have demonstrated that intracellular localization of *TAS2R38* in association with lipid droplets facilitates activation by bacterial quorum-sensing molecules, leading to MAPK signaling and upregulation of ABCB1 expression.^34^ These findings appear to contrast with our results and underscore that bitter receptor signaling is highly context-dependent. Rather than acting through uniform mechanisms, different T2Rs appear to have diverse roles, with some receptors repressing efflux and enhancing chemosensitivity, and others promoting chemoresistance.^33^ These observations suggest that the biologic consequences of T2R activation are likely determined by receptor subtype, subcellular localization, cellular context, and downstream signaling networks rather than by a conserved receptor program. In HNSCC, our results support a chemo-sensitizing role of caffeine, although whether this is mediated specifically though T2R10 remains to be established.

Although our findings parallel prior reports linking T2R10 activation to chemo-sensitization,^1^ we did not directly demonstrate that the effects of caffeine are exclusively mediated through T2R10. Caffeine has multiple biologic effects, including antagonism of adenosine receptors, inhibition of phosphodiesterases, and modulation of intracellular calcium flux.^35^ Our findings demonstrate that caffeine exposure increased *TAS2R10* promoter activity and receptor expression, raising the possibility that caffeine may modulate *TAS2R10*-associated pathways through transcriptional regulation in addition to previously described canonical receptor activation mechanisms. Definitive mechanistic validation will require targeted studies using CRISPR-based editing and selective agonists and antagonists. In addition, transporter expression was assessed at the mRNA level only, and future studies incorporating protein level and functional efflux assays will be important. Nonetheless, the correlation of high *TAS2R10* expression with favorable disease-free survival suggests that receptor-specific effects may be biologically relevant in HNSCC.

The caffeine concentrations used in this study, 100–200 μM, exceed the plasma levels typically achieved through routine dietary consumption, which may limit the direct translation of these findings to systemic caffeine administration.^35–38^ However, plasma concentrations may not accurately reflect transient local exposure of the oral and oropharyngeal epithelium, as caffeine concentrations in commonly consumed beverages exceed those used in our experiments. The duration and tissue penetration of such exposures are unknown, and our study did not evaluate whether comparable concentrations could be achieved or sustained within tumors in vivo. Nevertheless, these findings raise the possibility that caffeine or more selective T2R10-targeting compounds could be administered locally, such as through topical or intratumoral delivery, thereby reducing the need for high systemic exposure. Future studies should determine local pharmacokinetics, tissue penetration, safety, and therapeutic efficacy in physiologically relevant models. Overall, these findings establish a foundation for further investigation of TAS2R10-associated signaling and locally delivered receptor-targeted strategies as potential approaches to overcome chemoresistance in HNSCC.

## Acknowledgments

We thank M. Victoria (University of Pennsylvania) for excellent technical assistance and helpful discussions. This study was supported by the National Institutes of Health under Grants R01DE034474 (R.M.C.), R01HL178613 (R.J.L.), R01AI167971 (R.J.L.).

